# Exploring the potential role of the *TETRATRICOPEPTIDE THIOREDOXIN-LIKE* gene family in nitrogen-fixing and water-restricted soybean plants

**DOI:** 10.64898/2026.06.22.733792

**Authors:** Maria Martha Sainz, Carla Valeria Filippi, Selene Píriz-Pezzutto, Guillermo Eastman, José Sotelo Silveira, Omar Borsani, Mariana Sotelo Silveira

## Abstract

The TETRATRICOPEPTIDE THIOREDOXIN-LIKE (TTL) proteins are a plant-specific family proposed to function as peripheral membrane proteins that contribute to abiotic stress tolerance in Arabidopsis, likely by maintaining cell wall integrity through brassinosteroid signaling. Previously, we identified a *TTL* gene that was differentially regulated at the translational level in nitrogen-fixing soybean plants under water deficit (WD) conditions. This finding prompted the characterization of the soybean *TTL* gene family. Using the *Glycine max* v4.0 proteome, we identified ten TTL homologs (GmTTL1–GmTTL10), which are unevenly distributed across five chromosomes. Phylogenetic and structural analyses grouped these genes into three clades and revealed a highly conserved exon-intron organization. Likewise, GmTTL proteins display a conserved number and arrangement of TPR and TRXL motifs. To gain insights into their potential biological functions, we integrated co-expression and differential expression analyses. This approach identified a co-expression module enriched for translationally downregulated genes related to the Gene Ontology terms “cellular anatomical entity”, “membrane”, “cell periphery”, “cell wall modification”, “nitrate assimilation”, and “cell wall organization or biogenesis”. Protein-protein interaction network analysis of this specific subset of genes uncovered a novel GmTTL connection with two nitrate reductase enzymes in nitrogen-fixing plants subjected to WD, potentially linking the *TTL* gene family to new functions or roles. This study provides a framework for future functional studies of GmTTL proteins and their contribution to abiotic stress adaptation in soybean.

**Key Message:** This work presents the first functional characterization of TTLs proteins in legume species and highlights key processes that may link the *TTL* gene family to new functions or roles.

## 1. Introduction

The *TETRATRICOPEPTIDE THIOREDOXIN-LIKE (TTL)* gene family was first described in *Arabidopsis thaliana* (Arabidopsis). Their proteins are characterized by the presence of six tetratricopeptide repeats (TPR) in conserved positions and a non-functional thioredoxin-like domain in the carboxyl-terminal region, which shows homology to thioredoxins but lacks the essential cysteine residues required for thioredoxin activity (Lakhssassi et al., 2012; Rosado et al., 2006). The Arabidopsis *TTL* gene family comprises four members, with the first characterized member, *TTL1*, whose mutation causes root swelling and root growth arrest under NaCl and osmotic stress (Lakhssassi et al., 2012; Rosado et al., 2006). Furthermore, *TTL1* has been shown to contribute to anisotropic root growth during osmotic stress adaptation and to maintain cell wall integrity through a genetic interaction with *Cellulose synthase 6* (*CESA6)* (Cuadrado-Pedetti et al., 2021). Thanks to the presence of the TPR motifs, TTL proteins likely mediate the assembly of multiprotein complexes. Studies conducted with TTL3 showed that this protein interacts with a constitutively active BRASSINOSTEROID INSENSITIVE1 (BRI1) receptor kinase, BRI1-SUPPRESSOR1 phosphatase, and the BRASSINAZOLE RESISTANT1 transcription factor to optimize signal transduction in Arabidopsis (Amorim-Silva et al., 2019); TTL3 also interacts with microtubules and may link cytoskeletal function to the brassinosteroid signaling pathway (Xin et al., 2022). Additionally, TTL3 interacts with cellulose synthases, which modulate stress tolerance (Kesten et al., 2022). Moreover, the *TTL3* gene has been identified as being related to lateral root emergence and development (Xin et al., 2022).

This family of proteins is characteristic of land plants, as confirmed by extensive investigations across a variety of sequenced genomes. These proteins were not identified in animals or microorganisms but are present in all plant genomes analyzed, including those of *Physcomitrella patens* and *Selaginella moellendorffii.* Notably, they are absent in algae (Lakhssassi et al., 2012).

In recent studies conducted by our group, which aimed to analyze the differential responses of roots of nodulated (N-fix) soybean plants to water deficit (WD) at the transcriptional and translational levels, we identified a soybean *TTL* gene that is differentially expressed at the translational level (Sainz et al., 2024, 2022). Translational control has proven important in plants exposed to environmental stresses like WD (Kawaguchi et al., 2004; Lei et al., 2015), which is logical because this stage of gene expression regulation provides plants, and organisms in general, with flexibility and adaptability. It does not require new messenger RNA (mRNA) synthesis but instead depends on how efficiently existing mRNAs are translated (Lee and Bailey-Serres, 2019; Urquidi Camacho et al., 2020). Consequently, the analysis of the translatome—the subset of mRNAs that are being translated—enables more precise and comprehensive measurement of cell gene expression than analyzing only steady-state mRNA levels—the transcriptome (Sablok et al., 2017).

Cultivated soybean is a globally important crop that serves as the primary source of protein and oil for human and animal consumption and is increasingly becoming a key biodiesel crop (Vargas-almendra et al., 2024). Although there is extensive information regarding the response mechanisms of soybean to WD, most of these studies have been conducted on non-nodulated fertilized (N-fed) plants, even though producers mainly rely on the plant’s high capacity for symbiotic nitrogen fixation (Maluk et al., 2023). Therefore, analyzing the response of N-fix plants to WD appears to be more suitable, especially considering that it is known to differ from that of N-fed plants (Álvarez-Aragón et al., 2023; López et al., 2023; Sainz et al., 2024, 2022; Staudinger et al., 2016).

In this context, considering the previously characterized role of Arabidopsis *TTLs* mentioned before, we were interested in identifying the TTL family members and conducting an initial classification in soybean. Here, we identified ten TTL (Tetratricopeptide Thioredoxin-Like) homologs from the *Glycine max* v4.0 proteome. These homologs, which we named GmTTL1–GmTTL10, showed uneven physical locations, distributed across five chromosomes. The phylogenetic and structural studies performed showed their division into three clades and the conservation of their gene structure. The GmTTLs proteins showed conservation in the number and position of the TPR and TRXL motifs. The pairwise sequence comparison of the promoter regions of the *GmTTLs* showed 24% to 52% identity across all *GmTTLs.* The promoter region identity similarity correlated with the expression patterns of the *GmTTLs;* the higher the identity between two promoters, the more similar their expression patterns. Through gene co-expression and differential expression analysis, we identified a co-expression module enriched with down-regulated differentially expressed genes (DEGs) at the translational level in N-fix plants exposed to WD. One of these down-regulated DEGs was a *TTL* gene. By constructing a protein-protein interaction (PPI) network from this specific subset of DEGs, we uncovered novel, biologically meaningful associations regarding the role of this gene family in nodulated soybean roots subjected to WD. These findings provide a framework for future functional studies aimed at elucidating the contribution of GmTTL proteins to abiotic stress adaptation in soybean. Given that, to the best of our knowledge, GmTTL proteins have not yet been functionally characterized in legume species, this work establishes a valuable foundation for future experimental validation.

## 2. Materials and Methods

### 2.1. Retrieval of sequences and TTL family members identification and initial characterization

We retrieved sequences and annotation files from the *Glycine max* v4.0 reference proteome (GCF_000004515.6) from https://www.ncbi.nlm.nih.gov/ (The National Center of Biotechnology (NCBI). We used Interproscan v5.0 (Jones et al., 2014) with default parameters for protein domain annotation and domain prediction composition. Sequences containing at least one TPR and TRX domain were extracted, and BLASTP v2.13.0 (Altschul, 2014) was used to confirm them as GmTTLs defined and considered for posterior analysis.

Based on the protein sequence information, we used seqinr (Charif and Lobry, 2007) to compute isoelectric point and molecular weight (in kDa), while the prediction of subcellular localization of each protein was done using tPlant-mPLoc v2.0 (Chou and Shen, 2010). To determine the chromosomal location and gene structure of *GmTTL* homologs, we used the GFF3 annotation file. In addition, PlantCARE (accessed on 10 May 2022, (Lescot et al., 2002) was used to analyze the 2,000 bp genomic sequences located on the 5’ upstream of the Transcriptional Start Site (TSS) *GmTTLs* sequences to predict cis-regulatory elements (CRE) of *GmTTLs*. We analyzed the protein motifs using the MEME suite v5.4.1 (Bailey et al., 2015), with the maximum motif number set to 10 and an optimum width range set to 6-50 amino acids, the remaining parameters as default. We annotated the identified motifs using Interproscan v5.0 (Jones et al., 2014). To plot *GmTTLs* homologs gene structure and protein motif, added to CRE elements, we use R base functions (R Core Team, 2021) and pheatmap v1.0.12 (accessed on 1 June 2022; (Kolde, 2019). These upstream sequences were aligned, and pairwise nucleotide dissimilarity was calculated from the alignment as a measure of sequence distance. The resulting distance matrix was used for hierarchical clustering and visualized as a clustered heatmap in R using the pheatmap package (Kolde, 2025).

Furthermore, we investigated the transcript accumulation of the *GmTTLs* in soybean tissues from publicly available RNAseq data in Phytozome (https://phytozome.jgi.doe.gov/pz/portal.html) (Goodstein et al., 2012). Fragments Per Kilobase of transcript per Million mapped reads (FPKM) values were used to standardize gene expression levels across samples, accounting for gene length and sequencing depth. Afterward, the transcript accumulation pattern was compared with the promoter region identity similarity.

For comparison purposes, Arabidopsis TTL protein sequences were retrieved from The Arabidopsis Information Resource (TAIR, https://www.arabidopsis.org/), *Glycine max (GCF_000004515.6)*, *Lotus japonicus* (20210713_LjGifu1.3) and *Medicago truncatula* (GCF_003473485.1) from the NCBI (https://www.ncbi.nlm.nih.gov/). Arabidopsis, *G. max, L. japonicus* and *M. truncatula* amino acid sequences of the TTL protein family were aligned using ClustalW, as implemented in the R package msa v1.22.0 (Bodenhofer et al., 2015). The obtained multiple sequence alignments were used as input in phangorn v2.11.1 (Schliep, 2011) to construct the neighbor-joining phylogenetic trees.

### 2.2. Plant growth and drought assay

We grew plants of the Don Mario 6.8i (DM) soybean genotype under controlled conditions in a growth chamber, as detailed in Sainz et al., 2024, 2022. We sowed three seeds per pot and carefully analyzed the homogeneity of the seedlings to avoid any interference related to developmental phenotype. The *Bradyrhizobium elkanii* strain U1302 grown in liquid YEM-medium (Vincent, 1970) was used for the inoculated plants.

For the water deficit (WD) assay, the experimental unit was one pot with one plant. The experimental design was completely randomized, with 20 pots corresponding to four combined treatments and five biological replicates (n = 5). The four combined treatments were nodulated (N) water-restricted (WR) plants (N+WR), nodulated well-watered (WW) plants (N+WW), non-nodulated (NN) water-restricted plants (NN+WR), and non-nodulated well-watered plants (NN+WW).

The seedlings grew without water restriction for the first 19 days after sowing (V2-3 developmental stage), maintaining the substrate at field capacity with B&D-medium (Broughton and Dilworth, 1971) supplemented with KNO_3_ (0.5 mM and 5 mM final concentration for N and NN plants, respectively). Watering withdrawal to the WR plants began on day 20 (day 0 of the WD period), while we maintained the WW plants without water restriction throughout the assay. We measured the substrate water content daily and the stomatal conductance of all plants from day 20 till the end of the WD period, as described in Sainz et al., 2024. We established the end of the WD period individually for each WR plant when the stomatal conductance value reached 50% of the value obtained on day 0 of the WD period. At this moment, we harvested the roots of each WR plant and kept them at -80°C until polysomal fraction purification. We harvested the roots of the WW plants simultaneously with those of the WR plants and kept them at -80°C until the polysomal fraction was purified.

### 2.3. Polysomal fraction purification by sucrose cushion centrifugation

#### Preparation of cytoplasmic lysates

We performed all the steps at 4°C or on ice, and all equipment and materials were pre-chilled and RNase-free. We pulverized frozen roots in liquid nitrogen and homogenized 2 mL of packed tissue in 4 mL of Polysome Extraction Buffer described in Sainz et al., 2022, using a mortar and pestle. The homogenate was maintained at 4°C with gentle shaking for 15 min, then clarified by centrifugation at 16,000 g for 15 min. Then, we filtered the homogenate with cheesecloth and repeated the centrifugation step. 500 µL of the supernatant was reserved for total RNA isolation (TOTAL). The remaining supernatant (2 mL) was centrifuged through sucrose cushions to purify polysomal fractions.

#### Sucrose cushion centrifugation and polysome purification

We made two layers of sucrose cushions (4.5 mL of 12% and 4.5 mL of 33.5%) from a 2M sucrose stock solution, a 10x salts stock solution (400 mM Tris-HCl, pH 8.4, 200 mM KCl, 100 mM MgCl2) used at 1x, 50 μg ml^-1^ cycloheximide, and 50 μg ml^-1^ chloramphenicol. The 2 mL clarified cytosolic extract was loaded on the two-layer’s sucrose cushions and centrifuged in a Beckman L-100K class S ultracentrifuge (W40 Ti swinging bucket rotor) at 4 °C for 2 h at 35,000 rpm. 13.2 mL Ultra-Clear tubes were used (Beckman Coulter, United States, 344059). After centrifugation, the polysomal fraction was resuspended in 200 μL of Polysome Resuspension Buffer (PRB; 200 mM Tris-HCl, pH 9.0, 200 mM KCl, 25 mM EGTA, 35 mM MgCl2, 5 mM DTT, 50 μg ml^-1^ cycloheximide, 50 μg ml^-1^ chloramphenicol) by pipetting up and down several times. The resuspended polysomal pellet was maintained for 30 min at 4°C, and then regular RNA purification was performed to obtain the polysome-associated mRNA (PAR) fraction.

### 2.4. TOTAL and PAR RNA fractions extraction and transcriptome sequencing

We used 750 µL of TRizol LS reagent (Invitrogen, United States, 10296-028) to homogenize the resuspended polysomal pellet and the extract reserved for total RNA isolation. After homogenization, the samples were incubated for 5 min at room temperature (RT), followed by the addition of 200 µL of chloroform. The mixture was then vigorously shaken for 15 sec, incubated at RT for an additional 10 min, and centrifuged at 4°C and 12,000 g for 15 min to facilitate phase separation. We transferred 500 µL from the upper phase to a new tube containing 375 µL of cold isopropanol and 0.5 µL of RNase-free glycogen (Invitrogen, United States, 10814-010), and then incubated at 4°C for 10 min. The RNA precipitate was collected by centrifugation at 12,000 g for 15 min and washed with 1 mL of cold 75% ethanol. After centrifugation, the RNA pellet was air-dried, resuspended in 50 µL of RNase-free water, and incubated at 65°C for 5 min. RNA concentration and integrity were measured using an Agilent 2100 bioanalyzer (Agilent Technologies, Inc., United States). Samples with a RIN (RNA integrity number) greater than 7.0 and containing more than 1.0 µg were sent to Macrogen Inc. (South Korea) for library preparation and sequencing. TruSeq Stranded mRNA paired-end (PE) cDNA libraries were made and sequenced by the Illumina high-throughput sequencing platform. TOTAL and PAR samples from three biological replicates per combined treatment (N+WR; NN+WR; N+WW; NN+WW) were sent for analysis.

### 2.5. Data analysis

The analysis of the sequencing data on which this study relied was detailed in Sainz et al., 2024, and the data are available in the NCBI Sequence Read Archive (SRA): BioSample accessions: SAMN30227622-SAMN30227645, BioProject ID: PRJNA868178.

Therefore, we suggest checking that reference for a thorough description of the analysis pipeline. In this section, however, we provide a brief overview of the primary analysis methods.

#### Processing of sequencing data

We used FastQC v0.11.9 (https://www.bioinformatics.babraham.ac.uk/projects/fastqc/) to visually inspect the data quality of the Illumina sequencing. We trimmed the adapters and low-quality bases using Trimmomatic v0.39 (Bolger et al., 2014), retaining PE reads with an overall Phred quality score greater than 30 and a length greater than 80 bp for subsequent analysis.

We mapped the reads to the *Glycine max* v4.0 transcriptome (GCF_000004515.6, retrieved from https://www.ncbi.nlm.nih.gov/) using Salmon V0.12.0, in quasi-mapping mode (Patro et al., 2017), and we quantified the transcript abundance.

The R/BioConductor package Tximport (Soneson et al., 2016) was used to convert the estimated transcript-level abundances to gene-level expression abundances. Descriptive statistics were estimated using R base functions.

#### Differentially expressed gene (DEG) analysis

Differential expression analyses were performed using DESeq2 v3.15 (Love et al., 2014). Genes with |log2FC| > 1 and adjusted p-value (padj) < 0.05 were considered differentially expressed in our study. The generated gene lists were filtered to keep only differentially expressed TTLs. Plots were generated using R base functions and the R package ggplot2 version 3.3.6 (Wickham, 2016).

#### Co-expression analysis

Gene co-expression networks were constructed in R using the WGCNA package v1.71 (Langfelder and Horvath, 2008). DESeq2-normalized expression data were filtered to remove lowly expressed genes, defined as genes with fewer than 50 counts in more than 50% of the samples, and genes with low expression variability. The soft-thresholding power was selected with the pickSoftThreshold function and set to 17 to approximate scale-free topology. Modules were identified using the one-step network construction and module detection function, with maxBlockSize set to the total number of retained genes, TOMType = "signed", and mergeCutHeight = 0.20.

The resulting network was visualized in R using the network package, retaining edges with adjacency values ≥ 0.15. Modules were represented with distinct colors, and differentially expressed genes were alternatively highlighted according to their regulation status.

#### Protein-protein interaction (PPI) analysis

Gene lists analysis was performed using STRINGdb v11.5 (Jensen et al., 2009) with default parameters, using protein sequences in FASTA format as input. For visualization purposes, unconnected nodes were hidden.

## 3. Results

### 3.1 TTL family member identification and initial characterization in soybean

Soybean proteome-wide analysis led to the identification of 302 proteins that have TPR domains (Table S1); the majority of them (234) have 1 to 3 TPR motifs, whereas the rest display between 4 to 13 TPR TPR motifs (Table S1). Previously, we have identified a total of 125 soybean proteins containing Trx motifs (Sainz et al., 2022).

The TTL proteins constitute a family specific to land plants that were first described in Arabidopsis for having six TPR domains located in specific positions throughout the sequence and a thioredoxin-like (TRXL) motif located in the C-terminal end of the sequence (Ceserani et al., 2009; Prasad et al., 2010; Rosado et al., 2006), which lacks an essential Cysteine required for reducing activity (Ceserani et al., 2009; Rosado et al., 2006).

A total of 11 TTL homologs were identified from the *Glycine max v4.0* proteome. After discarding a protein codified by the same gene, a total of 10 TTL homologs remained for further analysis (Table S2). For subsequent analysis, the TTL proteins were sequentially numbered from GmTTL1 to GmTTL10, following the same order as their NCBI IDs. GmTTL homolog lengths ranged from 584 (GmTTL6) to 703 (GmTTL7) amino acids. Accordingly, their molecular weights ranged from 63,9 kDa (GmTTL6) to 78.7 kDa (GmTTL7). The predicted values for the isoelectric point (pI) varied from 8.8 (GmTTL2 and GmTTL5) to 9.5 (GmTTL8) (Table S3). Regarding subcellular localization, all GmTTL homologs are predicted to be localized in the nucleus, in contrast to the cytoplasmic localization reported in Arabidopsis for these proteins (Kesten et al., 2022; Lakhssassi et al., 2012).

By mapping the 10 *GmTTL* homologs on the *Glycine max* chromosomes, we observed that their physical locations are distributed across five chromosomes: *GmTTL1* on chromosome 4, *GmTTL2* and *GmTTL3* on chromosome 6, *GmTTL4*, *GmTTL9,* and *GmTTL10* on chromosome 12, *GmTTL5*, *GmTTL6,* and *GmTTL7* on chromosome 13, and *GmTTL8* on chromosome 14 (Table S2).

### 3.2 Phylogenetic, gene structure, and conserved motif analyses of the *GmTTL* gene family

Phylogenetic analysis revealed that *GmTTL* genes could be divided into three subgroups (Figure 1a). Subgroups G1 and G2 consisted of 4 *GmTTL* members, whereas subgroup G3 contained 2 *GmTTL* members. The GmTTLs proteins showed conservation in the number and position of the TPR and TRXL motifs (Figure 1b).

**Figure 1.**
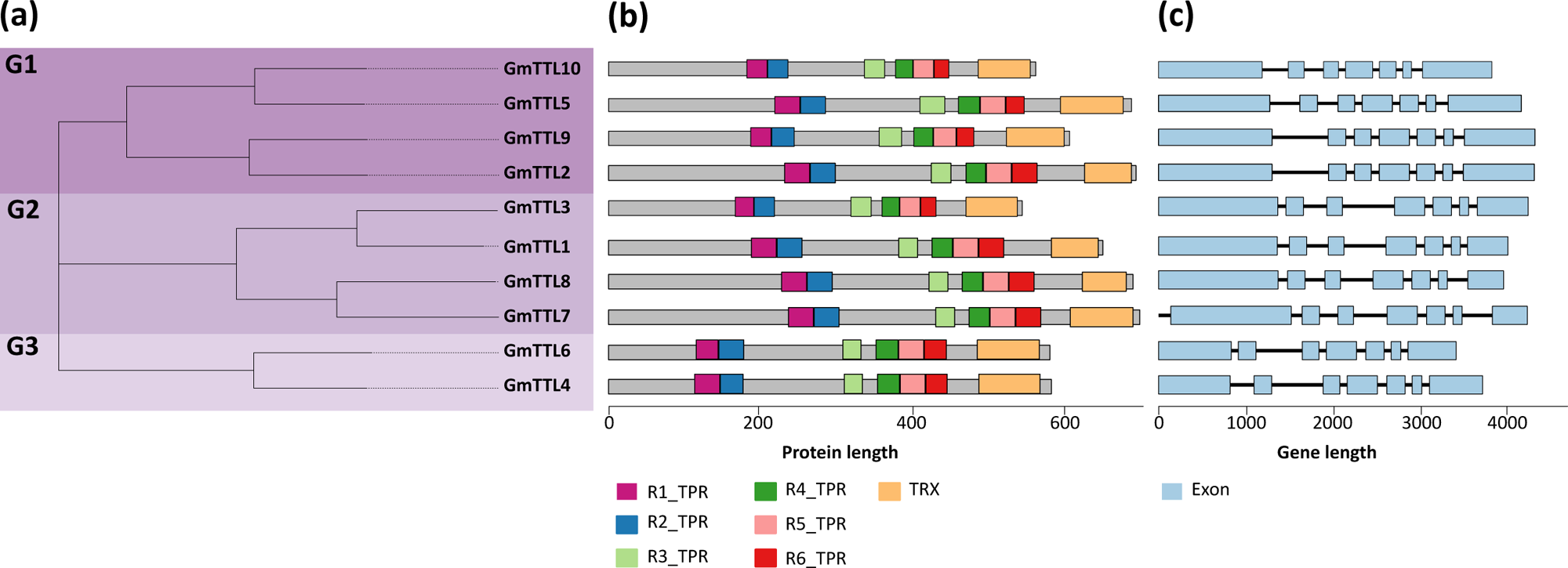
Phylogenetic tree, conserved motif, and gene structure of *Glycine max TTL* genes (*GmTTL*). (a) Phylogenetic relationship among the GmTTL based on the amino acid sequence alignment. (b) The GmTTLs proteins showed conservation in the number and position of the TPR and TRLX motifs. (c) Exon-intron structures of *GmTTL* genes.

The length of the *GmTTL* genes ranged from 3285 bp (*GmTTL6*) to 4156 bp (*GmTTL9*) (Supplementary Table S1b). The analysis of the exon-intron structures of *GmTTL* genes revealed that all members contained 7 exons (Figure 1c), ranging in size from 100 bp to 1330 bp. Curiously, the fourth exon is of the same length in all the genes of the family.

A phylogenetic analysis of all TTL proteins from *Arabidopsis thaliana*, *Lotus japonicus,* and *Medicago truncatula* also revealed that GmTTL proteins form three groups (Figure 2). The largest group (G1) included 8 of the 10 GmTTLs that grouped with AtTTL1 and AtTTL2 of Arabidopsis, and the majority of TTL proteins of *Lotus japonicus* and *Medicago truncatula*. G2 contained GmTTL3 and GmTTL1. Whereas G3 included only one TTL protein from *Lotus japonicus* LjGi1g1v0241600.1.

**Figure 2.**
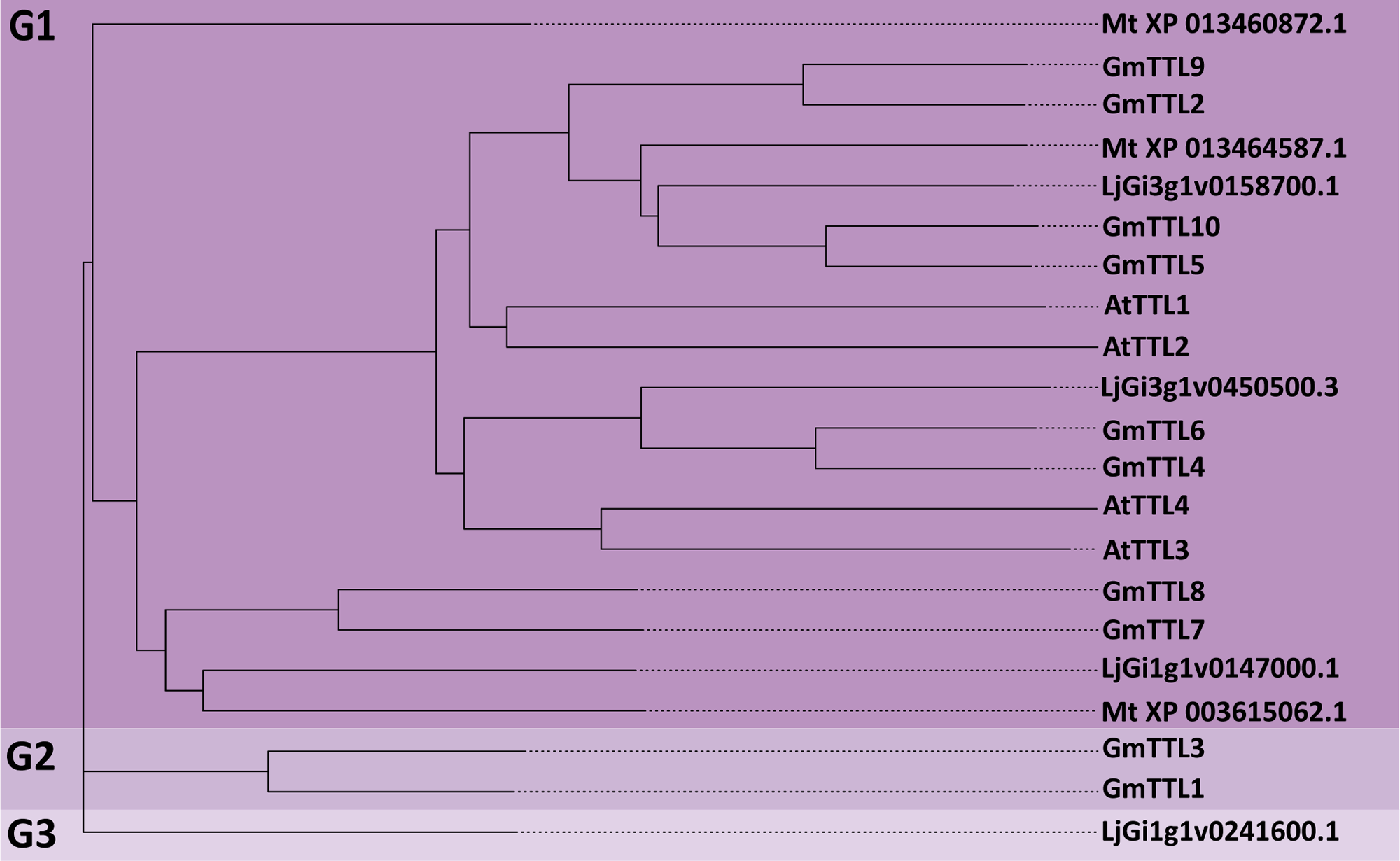
Phylogenetic tree of TTL proteins from *Glycine max*, *Medicago truncatula*, *Lotus japonicus and* Arabidopsis thaliana.

The phylogenetic analysis using the TRXL sequences showed the formation of three groups, G1, which includes all the AtTTLs together with six GmTTL proteins (GmTTL4, GmTTL6, GmTTL5, GmTTL10, GmTTL2, and GmTTL9), and G2, which includes 4 GmTTL (GmTTL1, GmTTL3, GmTTL7, and GmTTL8) that were more similar to the TTLs from *Medicago truncatula* and *Lotus japonicus*. The alignment of the TRXL domains showed that none of the GmTTL proteins contain the consensus sequence required for thioredoxin activity (Figure 3).

**Figure 3.**
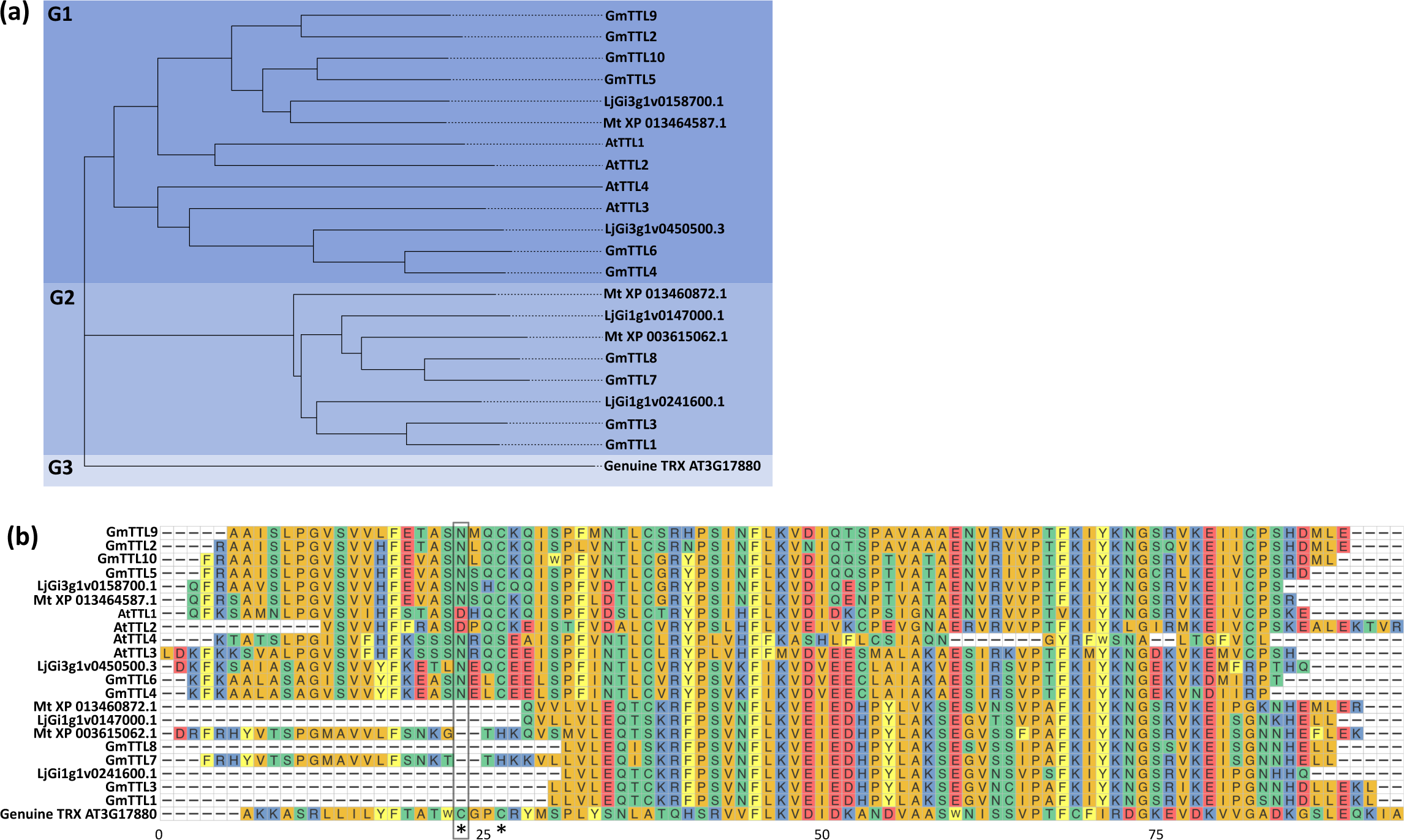
Phylogenetic and sequence analyses of TRXL domains. **(a)** Phylogenetic analysis of the TRXL domain of the TTLs from *Glycine max*, *Medicago truncatula*, *Lotus japonicus* and *Arabidopsis thaliana*. In the analysis, the protein sequence of At3g17880, a genuine thioredoxin, was included. **(b)** Details of the alignment of the TRXL domains, including the two-Cys motif (WCGPC) required for thioreductase activity. As shown, homologous regions from the TRXL domains of TTLs lack at least one of the two Cys residues that are essential for reductase activity.

As the TPRs motives in GmTTLs are located in similar positions in all the proteins, we performed the alignment of the six TPR motifs (R1_TPR to R6_TPR) between the proteins of Arabidopsis, soybean, *Medicago truncatula,* and *Lotus japonicus*. We confirmed that TPRs in the same position were more closely related to each other in other species than to other TPR motifs within the same GmTTL protein (Lakhssassi et al., 2012; Figure 4). Figure 4 shows how R1_TPR from GmTTL1 is more related to R1_TPR from *Lotus japonicus* than to other TPRs within GmTTL1.

**Figure 4.**
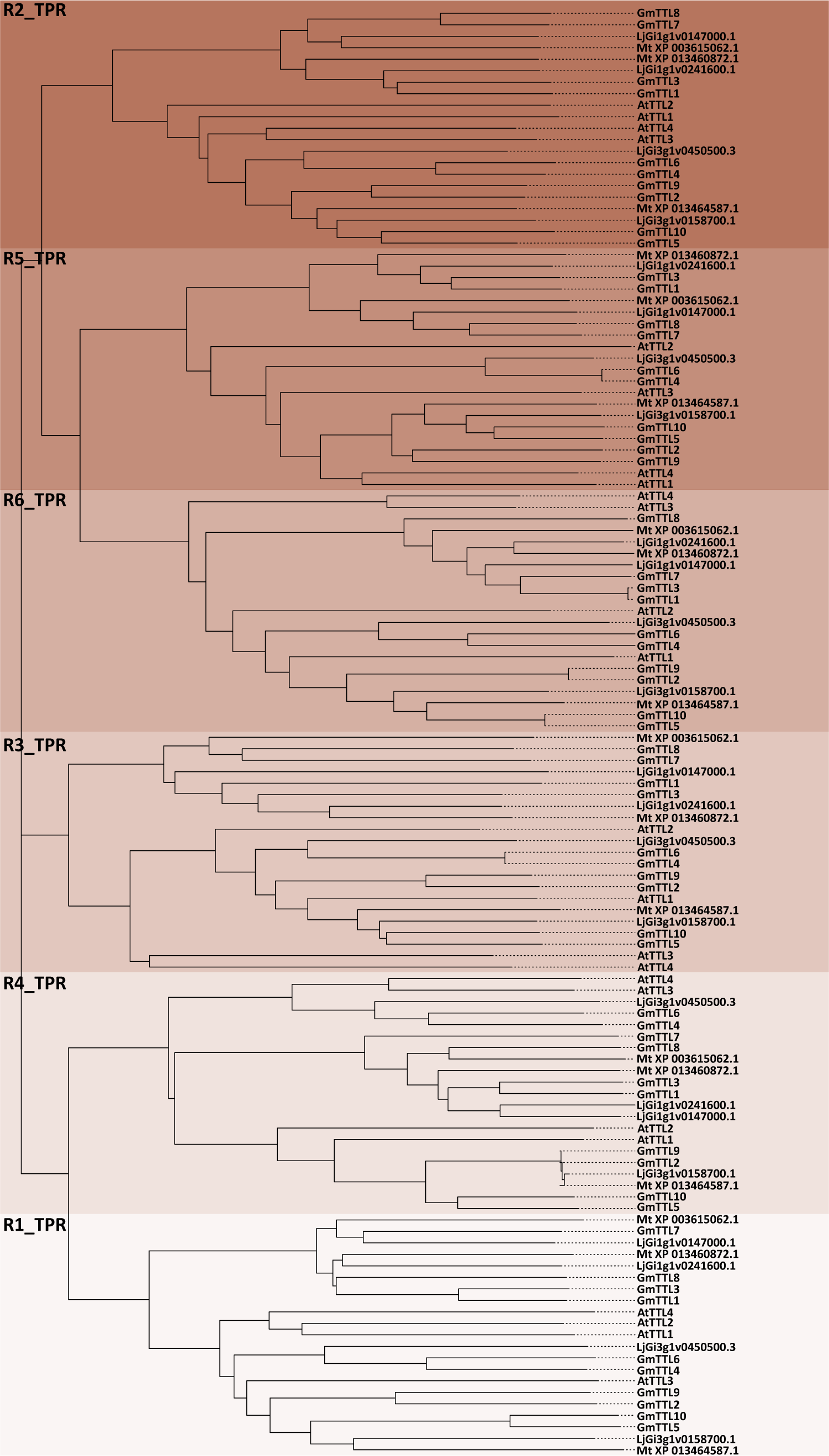
Phylogenetic relationship among individual TPR motifs. An unrooted tree was built using the six TPR sequences of the four *Glycine max* proteins and TPR sequences from TTL orthologs from *Medicago truncatula, Lotus japonicus*, and *Arabidopsis thaliana*.

### 3.3. Analysis of upstream regions and Cis-Regulatory Elements (CRE) for *GmTTL* and its expression patterns

We perform a pairwise sequence comparison between the promoter regions of the *GmTTLs* using 2000 bp upstream of the TSS. The analysis showed a range of 24% to 52% identity between the promoters of all *GmTTLs* (Figure 5a). The promoter sequences of *GmTTL1* and *GmTTL3* were the most similar (52% of identity; Figure 5a), and the promoter sequences of *GmTTL6* and *GmTTL10* were the least similar (24% of identity; Figure 5a). Interestingly, the majority of the promoters formed the same pairs as the protein sequences (Figure 1a); however, the promoters from *GmTTL2, GmTTL5, GmTTL9,* and *GmTTL10* formed different pairs (Figure 1a, Figure 5). We then investigated the transcript accumulation of the *GmTTLs* in soybean tissues from publicly available RNA-Seq data in Phytozome (https://phytozome.jgi.doe.gov/pz/portal.html) (Goodstein et al., 2012). The heatmap expression analysis of the entire family of *GmTTLs* genes (Figure S1) separated into two clusters, one that includes *GmTTL1*, *GmTTL3, GmTTL4, GmTTL6, GmTTL7, and GmTTL8; and the other that contains GmTTL2*, *GmTTL5*, *GmTTL9*, and *GmTTL10.* The first cluster is characterized by having genes with low levels of expression in seeds, leaves, shoots, root tip, and flowers. However, the second cluster is characterized by having genes with high expression in seeds, leaves, shoots, root tip, and flowers. Interestingly, some closely related genes based on the amino acid sequence showed similar expression patterns, as can clearly be seen for *GmTTL1* and *GmTTL3,* which both show high levels of expression in nodules, roots, and roots treated with urea, ammonium, and nitrate; but low levels in seeds, leaves, shoots, root tip, and flowers (Figure S1). The clustering of the expression pattern of *GmTTLs* genes resembled the grouping of the promoters by their sequence similarity (Figure 5, Figure S1).

**Figure 5.**
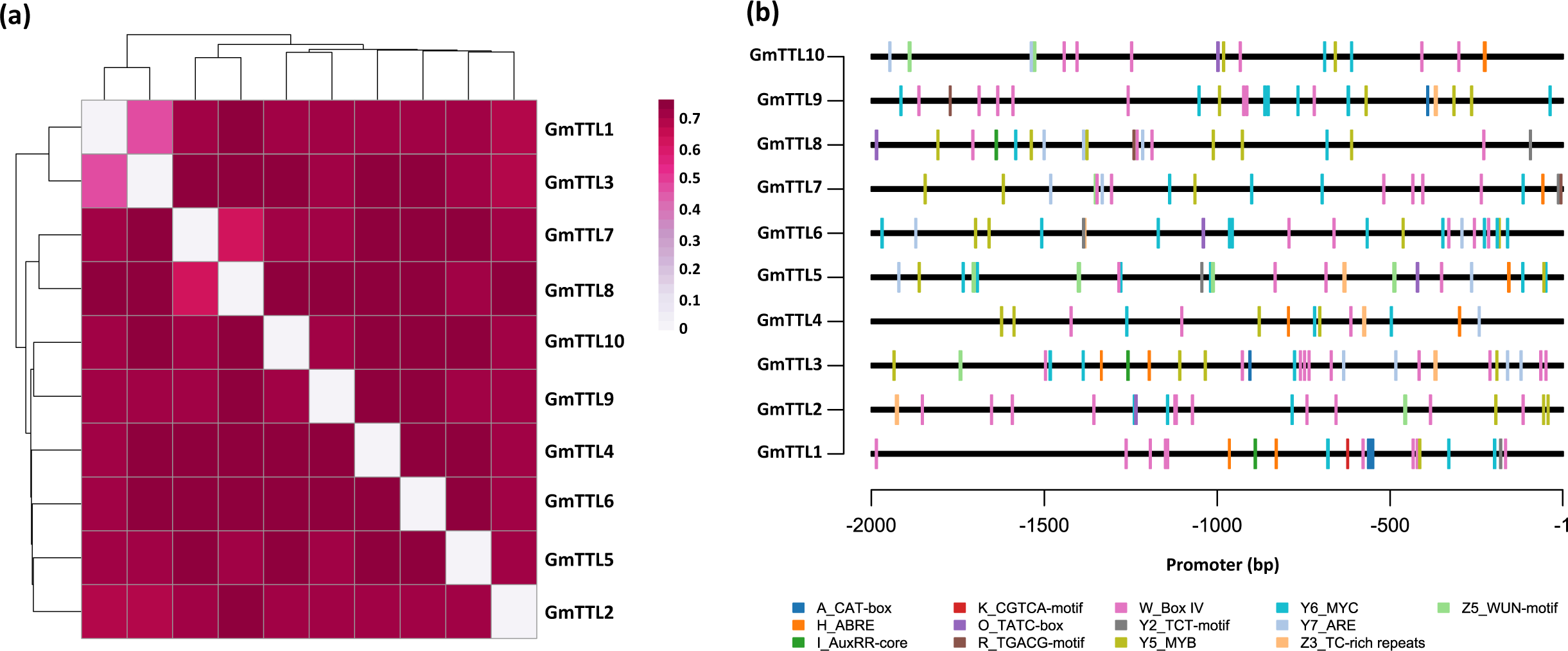
Analysis of promoter region and Cis-Regulatory Elements (CRE) for *GmTTL*. **(a)** Pairwise sequence comparison between the promoter regions of the *GmTTLs* using 2000 bp upstream of the TSS. The analysis showed a range of 24% to 52% identity between the promoters of all *GmTTLs.* **(b)** Number and localization of the more frequent cis-acting elements in the promoter regions of the *GmTTL* gene family. The 2000 bp genomic upstream of the Transcriptional Start Site (TSS) of *GmTTL* sequences were analyzed with PlantCARE (Lescot et al. 2002) to predict their cis-regulatory elements (CREs). The cis-acting elements presented are labeled in different colors and illustrated at the bottom. The TATA-box and CAAT-box are not shown.

To understand the potential regulatory mechanisms of *GmTTL* genes, we analyzed the presence of cis-regulatory elements 2000 bp upstream of the TSS. Based on their putative functions, the identified cis-acting elements were further classified according to the type of response (Figure 5b). Except for the common cis-acting elements (such as enhancer element CAAT-box and core promoter element TATA-box), the most abundant elements were stress-responsive elements, including WUN motifs that were present in 7 of 10 putative promoter regions, TC-rich repeats present in around 7 of 10 *GmTTL*, MYB binding site involved in drought inducibility (MBS) present in 10 of the *GmTTL’s* putative promoters, and the anaerobic induction ARE motif present in 8 of the 10 *GmTTL’s* putative promoters. Some CRE were involved in light responsiveness, such as Box IV, which was present in 100% of the *GmTTL* putative promoters. Moreover, we detected hormone-related cis-acting elements, including the MeJA-responsive element (TGACG motif and CGTCA motif), the ABA-responsive element (ABRE) present in 6 of 10 putative promoters, and the AuxRR core involved in auxin responsiveness was present in 3 of 10 putative promoter regions. Similar cis-regulatory elements were found in the *AtTTLs* promoter regions (Table S6) supported with experimental evidence (Cuadrado-Pedetti et al., 2021; Lakhssassi et al., 2012). This suggests that these genes are regulated by light, abiotic stresses, and hormones.

### 3.4 Identification of the differentially expressed *GmTTL* genes in N-fixing and water-restricted plants

The RNA-seq data—from TOTAL and PAR RNA fractions—used in this study were obtained from previous research in our lab, where a nodulation and WD experiment was conducted in soybean (Sainz et al., 2024, 2022). The experiment included four combined treatments: nodulated water-restricted plants (N+WR), nodulated well-watered plants (N+WW), non-nodulated water-restricted plants (NN+WR), and non-nodulated well-watered plants (NN+WW). These treatments allow for different comparisons. The two comparisons analyzed here highlight the differential response of N-fix and water-restricted plants: *i*) N+WR vs. N+WW and *ii*) N+WR vs. NN+WR. The first comparison illustrates the response of nodulated plants to WD, while the second highlights their distinct response compared to non-nodulated plants under the same conditions. Furthermore, because we analyzed both TOTAL and PAR RNA fractions, the plant responses to WD in the two possible nodulation conditions were further categorized based on their transcriptional, translational, or combined transcriptional and translational regulation.

We begin exploring the RNA-seq data by analyzing the differentially expressed genes (DEGs) in the two previously mentioned comparisons. The status of the DEGs (up- or down-regulated) and the regulation level (TOTAL, PAR, and TOTAL+PAR) where differential expression was identified are further distinguished (Figure 6a). Contrast *i* presented a larger absolute number of DEGs—about five times more—than contrast *ii*, indicating that the imposition of WD altered the expression of a greater number of DEGs than the nodulation condition of the plant. Additionally, more DEGs were down-regulated than up-regulated in contrast *i*, which aligns with previously published literature showing a general down-regulation of gene expression in plants subjected to WD. In most cases (all except down-regulated DEGs from contrast *ii*), the majority of DEGs were at the TOTAL+PAR level, indicating that most genes are regulated at both the transcriptional and translational levels. For contrast *ii*, however, analysis showed that while there were more down-regulated DEGs than up-regulated at the TOTAL and PAR levels (144/62 and 77/36, respectively), at the combined TOTAL+PAR regulation level, the majority of DEGs were up-regulated (242/123). All of this data is depicted in Figure 6a.

**Figure 6.**
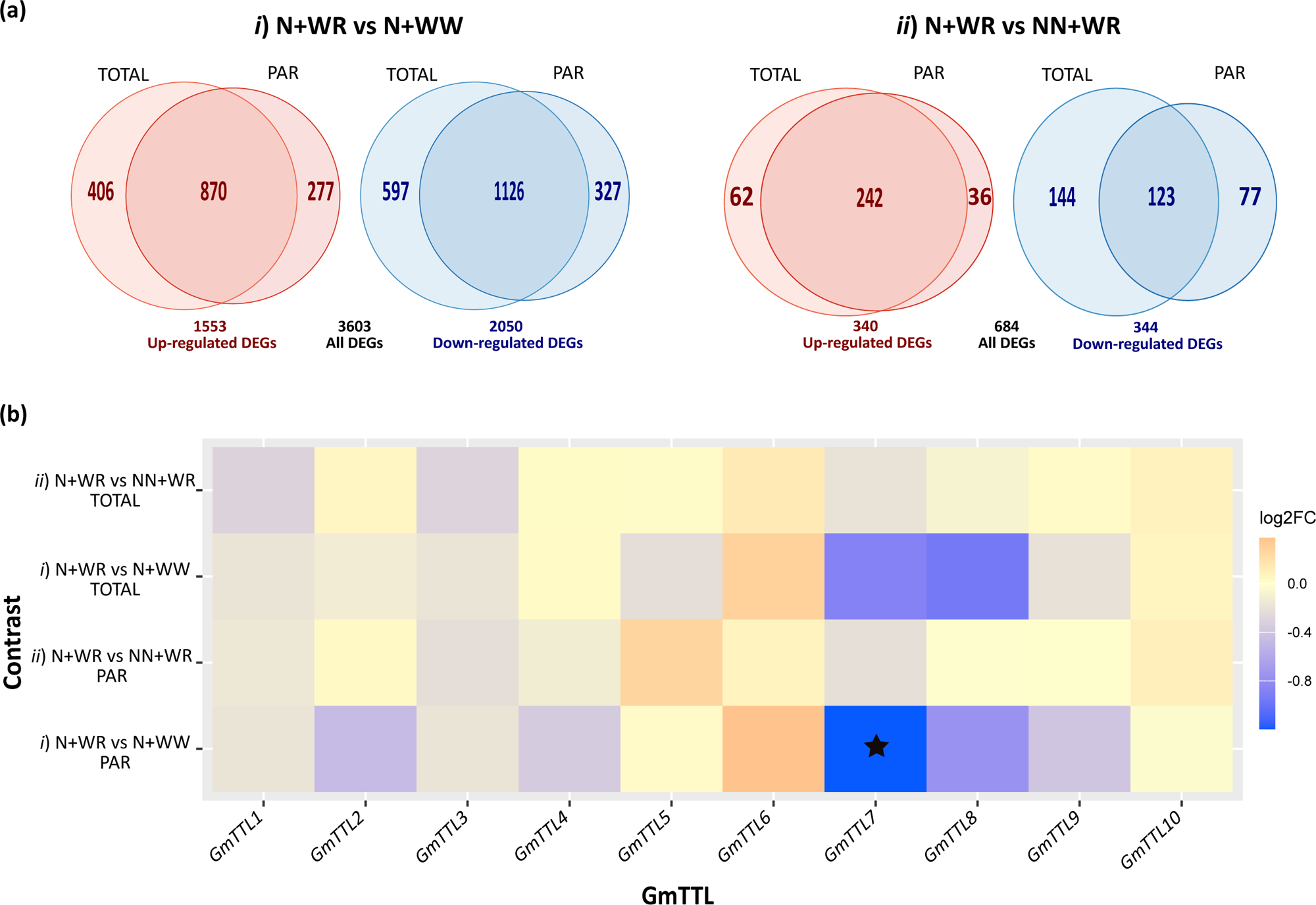
Differentially expressed gene (DEG) analysis and *GmTTL* expression profile in nodulated (N) and water-restricted (WR) plants. **(a)** Venn diagrams showing up- and down-regulated genes in the *i*) N+WR vs. N+WW and *ii*) N+WR vs. NN+WR contrasts in total RNA (TOTAL) and polysome-associated mRNA (PAR) fractions. **(b)** Expression profiles of *GmTTL*. The heatmap was constructed from the RNA-seq experimental data. An asterisk indicates the differentially expressed *GmTTL* gene found in our study. Genes with |log2FC| > 1 and adjusted p-value (padj) < 0.05 were considered differentially expressed.

Even though most DEGs were regulated at the transcriptional and translational levels (TOTAL+PAR), it is noteworthy that in both contrasts—encompassing both up- and down-regulated DEGs—some genes were regulated mainly at the translational level (PAR)(Figure 6a). This observation is consistent with the role of translational control of gene expression in enabling rapid cellular responses to stimuli, thereby providing greater flexibility and adaptability (Merchante et al., 2017).

Among these extensive DEG lists, we focused our analysis on the land plant-specific *GmTTL* genes to gain more insights into their role in nodulated plants exposed to environmental stresses. All ten *GmTTL* were detected in our RNA-seq data, and their expression levels in contrasts *i* and *ii*, as well as in the TOTAL and PAR fractions, are shown in Figure 6b. As expected, the expression profiles of the TOTAL and PAR fractions for each contrast showed a high correlation between them. Notably, *GmTTL7*—which presents high homology with *TTL3* from Arabidopsis (Figure 3a)—was the only gene exhibiting differential expression levels in *i*) N+WR vs N+WW at the translational (PAR) level. This is indicated with an asterisk in the corresponding profile of Figure 6b.

### 3.5 Downregulation of GmTTL7 in N-fixing plants under water restriction involves translational control

Another step in the exploration of the RNA-seq data was a weighted gene co-expression network analysis (WGCNA) followed by a differential expression analysis for identifying modules associated with the plant’s responses to the different treatments (Sainz et al., 2024) (Table S4). This type of analysis has proven useful for data interpretation, as it reduces the complexity of RNA-seq data and identifies modules that may reveal functional relationships between genes and their connection to biological processes (VanDam et al., 2018), such as those underlying the responses of N-fix plants to WD. Moreover, by combining co-expression and DEG analysis (Li et al., 2020; Sánchez-Baizán et al., 2022; Sferra et al., 2023) (i.e., DEGs in each fraction and comparisons between combined treatments were co-localized in the co-expression network modules), we focus our attention on module 7 (M7) because *GmTTL7*—the only *GmTTL* differentially expressed in nodulated and water-restricted plants—was localized there. Specifically, M7 was enriched in down-regulated DEGs. Since the differential expression of *GmTTL7* was achieved at the PAR level in contrast *i*, this subset of DEGs (i.e., M7 down-regulated DEGs from contrast *i* at the PAR level; Table S5) was further analyzed through a PPI-enriched network using the online STRING tool. This tool combines known and predicted relationships between proteins, covering both physical interactions and functional associations (Szklarczyk et al., 2021). Figure 7 shows that this particular subset of DEGs is enriched in the following GO terms: “cellular anatomical entity”, “membrane”, “cell periphery”, “cell wall modification”, “nitrate assimilation”, and “cell wall organization or biogenesis”. Notably, all nodes (proteins) in the network are included in the “cellular anatomical entity”. In particular, GmTTL7, which is also included in the “membrane” and “cell periphery” terms, showed connections with two HSP70-2 (Heat shock 70 kDa protein 2) and two nitrate reductase enzymes (NR and INR2, a constitutive and an inducible enzyme, respectively) (Figure 7). Additionally, TTL7 showed two other connections, though with lower confidence (i.e., thinner lines), to the epsilon subunit of the AP-4 complex and to an uncharacterized BTB/POZ and TAZ domain-containing protein (Figure 7). The AP-4 complex (Adaptor Protein Complex 4) forms a non-clathrin-associated coat on vesicles departing the trans-Golgi network and may be involved in targeting proteins from that network to the endosomal-lysosomal system (Law et al., 2022). The other connecting node, although uncharacterized, is a proposed peripheral membrane protein involved in the endomembrane system. These connections could be related to the already described behavior of AtTTLs as membrane proteins that modulate stress tolerance by safeguarding cellulose synthases (Kesten et al., 2022).

**Figure 7.**
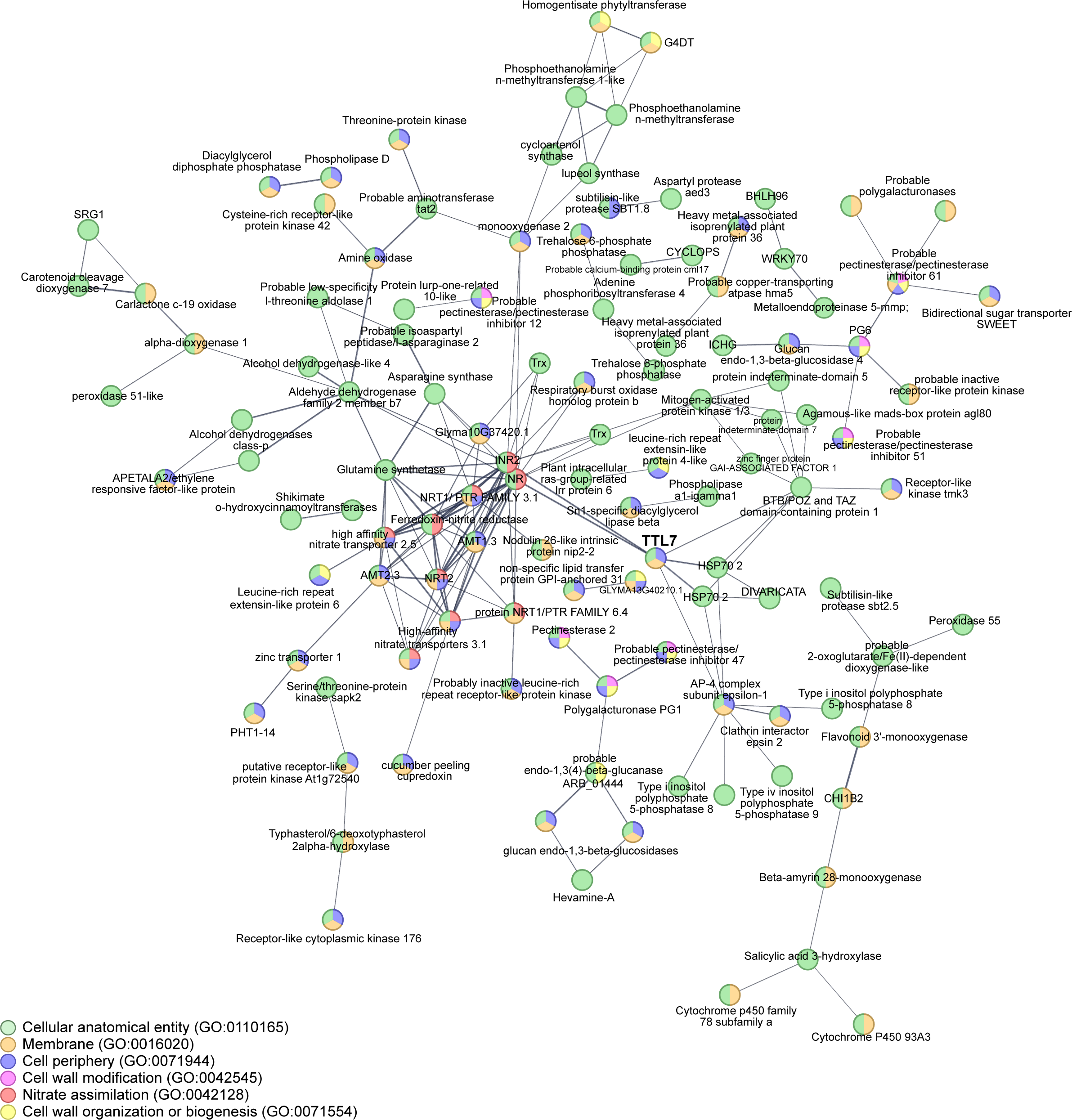
String analysis derived protein-protein interaction network obtained from WGCNA M7 down-regulated genes at the PAR level in nodulated and water-restricted plants with respect to nodulated and well-watered plants ((*i*) N + WR vs. N + WW comparison). This subset of differentially expressed genes is highlighted in bold in **Table S4**, and also is shown in **Table S5**. The network nodes represent proteins. Connecting lines denote protein-protein associations whose thickness indicates the strength of data support. The GO_BP terms are specified in the legends according to their colors. The number of nodes and edges are 356 and 171, respectively. PPI enrichment p-value < 1.0 x 10-16. Disconnected nodes, i.e., proteins that showed no interactions in the network, were removed.

The overrepresented functional terms identified in this PPI network highlight biological processes that may link the TTL gene family to new functions or roles.

## 4. Discussion

The *TETRATRICOPEPTIDE THIOREDOXIN-LIKE* (*TTL*) gene family is exclusive to land plants and was first described in *Arabidopsis thaliana* for its association with abiotic stress responses and developmental signaling (Amorim-Silva et al., 2019; Lakhssassi et al., 2012; Píriz-Pezzutto et al., 2024; Xin et al., 2022). However, despite growing knowledge of *TTL* function in Arabidopsis, this gene family remains largely unexplored in legume species and has not been characterized in *Glycine max*. In this study, we conducted the initial identification and description of the complete repertoire of the *TTL* family in soybean.

Based on the domain type related to protein function, we identified ten distinct *GmTTLs* genes in soybean (Table S2) that were distributed along five chromosomes. Overall, the phylogenetic analysis revealed that the *GmTTL* genes could be divided into three subgroups: G1 and G2, each with four members, and G3 with two members (Figure 1a). The structural features of the ten identified *GmTTL* genes resemble those originally described for Arabidopsis *TTLs* (Rosado et al., 2006). All members display strict conservation of exon numbers and sequence length conservation in the fourth exon across the entire family.

Furthermore, sequence analysis confirmed that the C-terminal thioredoxin-like (TRXL) domain of all 10 GmTTLs lacks the essential catalytic cysteine residues needed for disulfide reductase activity (Figure 3b). This result is consistent with findings in *Arabidopsis*, where TTL3 does not appear to possess reducing enzymatic activity, in agreement with its lack of conserved cysteine pairs found in active thioredoxins of the h class, yet is still considered important for protein stability (Ceserani et al., 2009). Interestingly, our phylogenetic analysis of TRXL domains separated the family into two distinct groups, with *GmTTL1/3/7/8* grouping more closely with legume TTLs orthologs than with Arabidopsis orthologs, suggesting a possible functional diversification specific to the Fabaceae family. The alignment of the six tetratricopeptide repeats (TPRs) demonstrated that individual TPRs are more closely related to their positional counterparts in other species (such as *Lotus japonicus* and *Medicago truncatula*) than to other TPR motifs within the same protein (Figure 4). This finding has already been described in Lakhssassi et al., 2012 and suggests that each TPR motif has evolved to facilitate highly specific, distinct protein-protein interactions within multiprotein bridging complexes, a mechanism vital for TTLs functioning as molecular scaffolds.

Interestingly, the pairwise sequence comparison between the promoter regions of the *GmTTLs* revealed a range of 24% to 52% identity between the promoters of all *GmTTLs* (Figure 5a), and the promoter sequences formed the same pairs as the protein sequences, except for promoters *GmTTL2, GmTTL5, GmTTL9,* and *GmTTL10* that formed different pairs (Figure 1a, Figure 5). Furthermore, we found a robust correlation between the sequence identity of the *GmTTL* promoter regions analyzed and their corresponding spatial expression patterns across tissues; the transcript accumulation of the *GmTTLs* in soybean tissues extracted from publicly available RNA-seq data separated two clusters, the first cluster is characterized by having genes with low levels of expression in seeds, leaves, shoots, root tip, and flowers and the second cluster characterized by having genes with high expression in seeds, leaves, shoots, root tip, and flowers. Notably, the clustering of GmTTL proteins according to amino acid sequence differed substantially from that based on expression patterns. However, some proteins, such as GmTTL1 and GmTTL3, showed concordance between both analyses, being closely related in sequence and exhibiting similar expression profiles. The clustering of the promoter regions also differs from the clustering observed with the GmTTL amino acid sequences, suggesting sub-functionalization driven by mutations in their regulatory sequences, as was observed with AtTTL2 (Lakhssassi et al., 2012).

The most abundant cis-regulatory elements found in the promoters of the *GmTTLs* were stress-responsive elements, including MYB-binding sites involved in drought inducibility, and those related to hormone induction, such as the ABA-responsive element (ABRE) and the AuxRR core involved in auxin responsiveness. Similar cis-regulatory elements were found in the *AtTTLs* promoters (Table S6) supported with experimental evidence (Cuadrado-Pedetti et al., 2021; Lakhssassi et al., 2012). This suggests that these genes are regulated by abiotic stresses and hormones in both species.

In this study, we provide initial insights into this gene family’s potential involvement in the response of soybean nitrogen-fixing plants subjected to water restriction. To achieve this, we leveraged RNA-seq datasets from a previous study that analyzed transcriptional and translational regulation of nitrogen-fixing and water-restricted soybean plants (Sainz et al., 2024). All ten *GmTTL* were detected in our RNA-seq data. Notably, the only gene exhibiting differential regulation—specifically downregulation—at the PAR level in N-fix and water-restricted plants was *GmTTL7*, which presents high homology to *TTL3 and TTL1* from Arabidopsis (Figure 3a) (Cuadrado-Pedetti et al., 2021; Rosado et al., 2006). To infer the functional context of this selective downregulation, we performed a co-expression analysis (VanDam et al., 2018) in combination with DEG analysis (Li et al., 2020; Sainz et al., 2024; Sánchez-Baizán et al., 2022; Sferra et al., 2023) and then constructed a protein-protein interaction network using STRING (Szklarczyk et al., 2021) with the WGCNA M7 subset of down-regulated genes from contrast *i* at the PAR level that includes *GmTTL7*. The overrepresentation of Gene Ontology (GO) terms such as "cellular anatomical entity", "membrane", "cell periphery", "cell wall organization or biogenesis", and "nitrate assimilation" provides valuable clues regarding the cellular processes potentially associated with *GmTTL7*. In Arabidopsis, TTL proteins have been proposed to function as bridges linking stress perception to the dynamic regulation of cellulose biosynthesis at the plasma membrane (Kesten et al., 2022). Although experimental validation is required, the enrichment of membrane- and cell wall-related functions in the present study is consistent with the possibility that GmTTL7 participates in related processes in soybean nodulated roots under WD.

Here, we found that GmTTL7 is associated with key components of cellular stress response and membrane trafficking, including HSP70-2 and the epsilon subunit of the AP-4 complex, which participates in vesicle trafficking from the trans-Golgi network to the endosomal-lysosomal system (Law et al., 2022). These associations, together with evidence of stress-induced relocalization of TTL3 to the plasma membrane in Arabidopsis (Kesten et al., 2022), support the hypothesis that GmTTL7 may be involved in membrane-associated processes contributing to cellular adaptation under water deficit.

Perhaps the more novel finding was the association of GmTTL7 with proteins involved in nitrate uptake and reduction. Combined with the reported role of AtTTL3 in lateral root emergence and development (Xin et al., 2022), these findings suggest that GmTTL7 may participate in a sensory or regulatory hub that coordinates root cell wall status, root growth, and metabolic shifts associated with nitrogen management during WD (Dziewit et al., 2026).

The translationally regulated network identified in this study provides a set of candidate genes and interactions for future experimental validation. Functional approaches such as gene knockout or knockdown strategies, protein interaction assays, and subcellular localization analyses will be valuable for elucidating the role of GmTTL7 in the adaptation of soybean roots to water deficit.

## Supplementary material

### Figures

**Figure S1.** Heatmap showing the expression analysis of the entire family of *GmTTLs* genes. Analysis of the transcript accumulation of the *GmTTLs* in soybean tissues from publicly available RNA-Seq data in Phytozome (https://phytozome.jgi.doe.gov/pz/portal.html) (Goodstein et al., 2012).

## Tables

**Table S1. Predicted TPR repeat regions in *Glycine max* proteins.**

Tetratricopeptide repeat (TPR) motifs predicted in proteins of *Glycine max* (soybean), including the repeat number (ordered from N-terminus to C-terminus), the start and end amino acid positions within each protein, and their corresponding NCBI Protein IDs. Predictions were obtained using InterProScan.

**Table S2. Genomic features and functional annotations of *TTL-like* genes in *Glycine max*.**

This table summarizes ten *Glycine max* (soybean) genes belonging to the TTL (TPR repeat-containing thioredoxin-like) protein family. For each gene, the table includes a short gene name (gene_name), stable gene identifier (stable_id), NCBI gene ID (ncbi_gene_id), and functional annotation (annotation). Genomic coordinates are given by the chromosome (chromosome), gene start and end positions (gene_start, gene_end), and coding strand (strand). The corresponding predicted protein is indicated with its identifier (protein_id) and length in amino acids (protein_length). All gene models are based on version 4 of the soybean reference genome.

**Table S3. Isoelectric points, molecular weights, and predicted nuclear localization of TTL proteins in *Glycine max*.**

This table summarizes predicted biochemical properties of TTL proteins (TPR repeat-containing thioredoxin-like) from Glycine max. The data includes the gene short name (Gm_TTL_name), predicted isoelectric point (isoelectric_point), estimated molecular weight in kilodaltons (molecular_weight_kDa), and subcellular localization (predicted_localization). All proteins were predicted to localize to the nucleus.

**Table S4. Differentially expressed genes (DEGs), and their WGCNA module correspondence, in the *i*) N+WR vs N+WW contrast within the polysomal fraction.**

List of DEGs identified in the N+WR (nodulated + water restricted) vs N+WW (nodulated + well-watered) contrast, within the polysomal (P) mRNA fraction. Only genes meeting the significance thresholds (adjusted p-value < 0.05 and |log₂FoldChange| ≥ 1) are shown. Gene IDs correspond to NCBI Gene identifiers. GmTTL7, found to be differentially expressed in this contrast, is highlighted in color in the table. A. The product of these genes were used for protein-protein interaction analysis using the STRING database.

Module_WGCNA pertenence were defined in Sainz et al (2024), https://doi.org/10.1186/s12870-024-05280-5.

In bold, genes included in the protein-protein interaction analysis.

**Table S5. Genes included in the protein-protein interaction analysis.**

All genes belonging to Module 7 - Downregulated in the condition N+WR vs N+WW - Only PAR fraction.

**Table S6. Number and localization of the more frequent cis-acting elements among the *AtTTL* promoter regions.** The 2000 bp of genomic sequence upstream of the Transcriptional Start Site (TSS) of *AtTTL* sequences were analyzed with PlantCARE (Lescot et al., 2002) to predict their cis-regulatory elements (CREs).

## Data availability

All datasets supporting the results of this study are included within the article and its supplementary information. The sequencing data is available in the NCBI Sequence Read Archive (SRA) under the accession number PRJNA868178.

## Supporting information

Figure S1_Sainz et al.

Supplementary tables_Sainz et al.

## Acknowledgments

We thank the Sistema Nacional de Investigadores (ANII) (María Martha Sainz, Carla Valeria Filippi, Guillermo Eastman, José Sotelo-Silveira, Omar Borsani, Mariana Sotelo-Silveira).

## Author Contributions

Conceptualization, M.M.S. and M.S.-S.; Data curation, C.V.F., G.E. and J.S.-S.; Formal analysis, M.M.S., C.V.F., J.S.-S. and M.S.-S.; Funding acquisition, M.M.S. and O.B.; Investigation, M.M.S., S.P-P; Methodology, M.M.S. and J.S.-S.; Resources, J.S.-S.; Writing—original draft, M.M.S. and M.S.-S.; Writing—review and editing, M.M.S., C.V.F., G.E., J.S.-S., O.B. and M.S.-S. All authors have read and agreed to the published version of the manuscript.

## References

1. Altschul, S.F., 2014. BLAST Algorithm . eLS 1–5. 10.1002/9780470015902.a0005253.pub2

2. Álvarez-Aragón, R., Palacios, J.M., Ramírez-Parra, E., 2023. Rhizobial symbiosis promotes drought tolerance in Vicia sativa and Pisum sativum. Environ. Exp. Bot. 208. 10.1016/j.envexpbot.2023.105268

3. Amorim-Silva, V., Garcia-Moreno, A., Castillo, A.G., Lakhssassi, N., Esteban del Valle, A., Perez-Sancho, J., Li, Y., Pose, D., Perez-Rodriguez, J., Lin, J., Valpuesta, V., Borsani, O., Zipfel, C., Macho, A.P., Botella, M.A., 2019. TTL proteins scaffold brassinosteroid signaling components at the plasma membrane to optimize signal transduction in Arabidopsis. Plant Cell tpc.00150.2019. 10.1105/tpc.19.00150

4. Bailey, T.L., Johnson, J., Grant, C.E., Noble, W.S., 2015. The MEME Suite. Nucleic Acids Res. 43, W39–W49. 10.1093/nar/gkv416

5. Bodenhofer, U., Bonatesta, E., Horejs-Kainrath, C., Hochreiter, S., 2015. msa: an R package for multiple sequence alignment. Bioinformatics 31, 3997–3999.

6. Bolger, A.M., Lohse, M., Usadel, B., 2014. Trimmomatic: a flexible trimmer for Illumina sequence data. Bioinformatics 30, 2114–2120. 10.1093/bioinformatics/btu170

7. Broughton, W.J., Dilworth, M.J., 1971. Control of Leghaemoglobin Synthesis in Snake Beans. Biochem. J. 125, 1075–1080. 10.1042/bj1251075

8. Ceserani, T., Trofka, A., Gandotra, N., Nelson, T., 2009. VH1 / BRL2 receptor-like kinase interacts with vascular-specific adaptor proteins VIT and VIK to influence leaf venation. Plant J. 1000–1014. 10.1111/j.1365-313X.2008.03742.x

9. Charif, D., Lobry, J.R., 2007. SeqinR 1.0-2: A Contributed Package to the R Project for Statistical Computing Devoted to Biological Sequences Retrieval and Analysis, in: In: Bastolla, U., Porto, M., Roman, H.E., Vendruscolo, M. (Eds) Structural Approaches to Sequence Evolution.

10. Chou, K.C., Shen, H. Bin, 2010. Plant-mPLoc: A top-down strategy to augment the power for predicting plant protein subcellular localization. PLoS One 5. 10.1371/journal.pone.0011335

11. Cuadrado-Pedetti, M.B., Rauschert, I., Sainz, M.M., Amorim-Silva, V., Botella, M.A., Borsani, O., Sotelo-Silveira, M., 2021. The Arabidopsis TETRATRICOPEPTIDE THIOREDOXIN-LIKE 1 Is Involved in Anisotropic Root Growth during Osmotic Stress Adaptation. Gene 12.

12. Dziewit, K., Wabnik, K., Szal, B., Podgórska, A., 2026. Nitrogen sources modulate auxin transport to fine - tune root system architecture. Plant Cell Rep. 10.1007/s00299-026-03849-y

13. Goodstein, D.M., Shu, S., Howson, R., Neupane, R., Hayes, R.D., Fazo, J., Mitros, T., Dirks, W., Hellsten, U., Putnam, N., Rokhsar, D.S., 2012. Phytozome : a comparative platform for green plant genomics. Nucleic Acids Res. 40, 1178–1186. 10.1093/nar/gkr944

14. Jensen, L.J., Kuhn, M., Stark, M., Chaffron, S., Creevey, C., Muller, J., Doerks, T., Julien, P., Roth, A., Simonovic, M., Bork, P., von Mering, C., 2009. STRING 8 - A global view on proteins and their functional interactions in 630 organisms. Nucleic Acids Res. 37, 412–416. 10.1093/nar/gkn760

15. Jones, P., Binns, D., Chang, H.-Y., Fraser, M., Li, W., McAnulla, C., McWilliam, H., Maslen, J., Mitchell, A., Nuka, G., Pesseat, S., Quinn, A., Sangador-Vegas, A., Scheremetjew, M., Yong, S.-Y., Lopez, R., Hunter, S., 2014. InterProScan 5: genome-scale protein function classification. Bioinformatics 30, 1236–1240.

16. Kawaguchi, R., Girke, T., Bray, E.A., Bailey-Serres, J., 2004. Differential mRNA translation contributes to gene regulation under non-stress and dehydration stress conditions in Arabidopsis thaliana. Plant J. 38, 823–39. 10.1111/j.1365-313X.2004.02090.x

17. Kesten, C., García-moreno, Á., Amorim-silva, V., Menna, A., Castillo, A.G., Percio, F., Armengot, L., Ruiz-lopez, N., Jaillais, Y., Sánchez-rodríguez, C., Botella, M.A., 2022. Peripheral membrane proteins modulate stress tolerance by safeguarding cellulose synthases. Sci. Adv. 6971, 1–13. 10.1126/sciadv.abq6971

18. Kolde, R., 2019. pheatmap: Pretty Heatmaps.

19. Lakhssassi, N., Doblas, V.G., Rosado, A., del Valle, A.E., Pose, D., Jimenez, A.J., Castillo, A.G., Valpuesta, V., Borsani, O., Botella, M.A., 2012. The Arabidopsis TETRATRICOPEPTIDE THIOREDOXIN-LIKE Gene Family Is Required for Osmotic Stress Tolerance and Male Sporogenesis. Plant Physiol. 158, 1252–1266. 10.1104/pp.111.188920

20. Langfelder, P., Horvath, S., 2008. WGCNA: An R package for weighted correlation network analysis. BMC Bioinformatics 9, 1–13. 10.1186/1471-2105-9-559

21. Law, K.C., Chung, K.K., Zhuang, X., 2022. An Update on Coat Protein Complexes for Vesicle Formation in Plant Post-Golgi Trafficking. Fontiers Plant Sci. 13, 1–9. 10.3389/fpls.2022.826007

22. Lee, T.A., Bailey-Serres, J., 2019. Integrative Analysis from the Epigenome to Translatome Uncovers Patterns of Dominant Nuclear Regulation during Transient Stress. Plant Cell 31, 2573–2595. 10.1105/tpc.19.00463

23. Lei, L., Shi, J., Chen, J., Zhang, M., Sun, S., Xie, S., Li, X., Zeng, B., Peng, L., Hauck, A., Zhao, H., Song, W., Fan, Z., Lai, J., 2015. Ribosome profiling reveals dynamic translational landscape in maize seedlings under drought stress. Plant J. 84, 1206–1218. 10.1111/tpj.13073

24. Lescot, M., Déhais, P., Thijs, G., Marchal, K., Moreau, Y., Peer, Y. Van De, Rouzé, P., Rombauts, S., 2002. PlantCARE , a database of plant cis-acting regulatory elements and a portal to tools for in silico analysis of promoter sequences. Nucleic Acids Res 30, 325–327.

25. Li, W., Wang, L., Wu, Y.U.E., Yuan, Z., Zhou, J., 2020. Weighted gene co - expression network analysis to identify key modules and hub genes associated with atrial fibrillation. Int. J. Mol. Med. 45, 401–416. 10.3892/ijmm.2019.4416

26. López, C.M., Alseekh, S., Torralbo, F., Martínez Rivas, Fc.D.S.J., Fernie, A.R., Amil-Ruiz, F., Alamillo, J.M., 2023. Transcriptomic and metabolomic analysis reveals that symbiotic nitrogen fixation enhances drought resistance in common bean. J. Exp. Bot. 74, 3203–3219. 10.1093/jxb/erad083

27. Love, M.I., Huber, W., Anders, S., 2014. Moderated estimation of fold change and dispersion for RNA-seq data with DESeq2. Genome Biol. 15, 1–21. 10.1186/s13059-014-0550-8

28. Maluk, M., Giles, M., Wardell, G.E., Akramin, A.T., Ferrando-molina, F., Murdoch, A., Barros, M., Beukes, C., Harrison, E., Daniell, T.J., Quilliam, R.S., Iannetta, P.P.M., James, E.K., 2023. Biological nitrogen fi xation by soybean ( Glycine max [ L .] Merr .), a novel , high protein crop in Scotland , requires inoculation with non-native bradyrhizobia. Front. Agron. 10.3389/fagro.2023.1196873

29. Merchante, C., Stepanova, A.N., Alonso, J.M., 2017. Translation regulation in plants: An interesting past, an exciting present and a promising future. Plant J. 90, 628–653. 10.1111/tpj.13520

30. Patro, R., Duggal, G., Love, M.I., Irizarry, R.A., Kingsford, C., 2017. Salmon provides fast and bias-aware quantification of transcript expression. Nat. Methods 14, 417–419. 10.1038/nmeth.4197

31. Píriz-Pezzutto, S., Martínez-Moré, M., Sainz, M.M., Borsani, O., Sotelo-Silveira, M., 2024. Arabidopsis root apical meristem adaptation to an osmotic gradient condition : an integrated approach from cell expansion to gene expression. Front. Plant Sci. 1–14. 10.3389/fpls.2024.1465219

32. Prasad, B.D., Goel, S., Krishna, P., 2010. In Silico Identification of Carboxylate Clamp Type Tetratricopeptide Repeat Proteins in Arabidopsis and Rice As Putative Co-Chaperones of Hsp90 / Hsp70. PLoS One 5. 10.1371/journal.pone.0012761

33. R Core Team, 2021. R: A laguage and environment for statistical computing.

34. Rosado, A., Schapire, A.L., Bressan, R.A., Harfouche, A.L., Hasegawa, P.M., Valpuesta, V., Botella, M.A., 2006. The Arabidopsis Tetratricopeptide Repeat-Containing Protein TTL1 Is Required for Osmotic Stress Responses and Abscisic Acid Sensitivity. Plant Physiol. 142, 1113–1126. 10.1104/pp.106.085191

35. Sablok, G., Powell, J.J., Kazan, K., 2017. Emerging Roles and Landscape of Translating mRNAs in Plants. Front. Plant Sci. 8, 1–9. 10.3389/fpls.2017.01443

36. Sainz, M.M., Filippi, C.V., Eastman, G., Sotelo-Silveira, J.R., Borsani, O., Sotelo-Silveira, M., 2022. Analysis of Thioredoxins and Glutaredoxins in Soybean: Evidence of Translational Regulation under Water Restriction. Antioxidants 11, 1–22.

37. Sainz, M.M., Filippi, C. V, Eastman, G., Sotelo-silveira, M., Zardo, S., Martínez-Moré, M., Sotelo-Silveira, J.R., Borsani, O., 2024. Water deficit response in nodulated soybean roots: a comprehensive transcriptome and translatome network analysis. BMC Plant Biol. 24, 1–18.

38. Sánchez-Baizán, N., Ribas, L., Piferrer, F., 2022. Improved biomarker discovery through a plot twist in transcriptomic data analysis. BMC Biol. 20, 1–26. 10.1186/s12915-022-01398-w

39. Schliep, K.P., 2011. phangorn: Phylogenetic analysis in R. Bioinformatics 27, 592–593. 10.1093/bioinformatics/btq706

40. Sferra, G., Fantozzi, D., Scippa, G.S., Trupiano, D., 2023. Key Pathways and Genes of Arabidopsis thaliana and Arabidopsis halleri Roots under Cadmium Stress Responses : Differences and Similarities. Plants 12, 1–19. 10.3390/plants12091793

41. Soneson, C., Love, M.I., Robinson, M.D., 2016. Differential analyses for RNA-seq: Transcript-level estimates improve gene-level inferences. F1000Research 4, 1–23. 10.12688/F1000RESEARCH.7563.2

42. Staudinger, C., Mehmeti-Tershani, V., Gil-Quintana, E., Gonzalez, E.M., Hofhansl, F., Bachmann, G., Wienkoop, S., 2016. Evidence for a rhizobia-induced drought stress response strategy in Medicago truncatula. J. Proteomics 136, 202–213. 10.1016/j.jprot.2016.01.006

43. Szklarczyk, D., Gable, A.L., Nastou, K.C., Lyon, D., Kirsch, R., Pyysalo, S., Doncheva, N.T., Legeay, M., Fang, T., Bork, P., Jensen, L.J., von Mering, C., 2021. The STRING database in 2021: customizable protein–protein networks, and functional characterization of user-uploaded gene/measurement sets. Nucleic Acids Res. 49, 10800. 10.1093/nar/gkab835

44. Urquidi Camacho, R.A., Lokdarshi, A., von Arnim, A.G., 2020. Translational gene regulation in plants: A green new deal. Wiley Interdiscip. Rev. RNA 11, 1–40. 10.1002/wrna.1597

45. VanDam, S., Võsa, U., van der Graaf, A., Franke, L., de Magalhães, J.P., 2018. Gene co-expression analysis for functional classification and gene-disease predictions. Brief. Bioinform. 19, 575–592. 10.1093/bib/bbw139

46. Vargas-almendra, A., Ruiz-medrano, R., N, L.A., Calder, B., Xoconostle-c, B., 2024. Advances in Soybean Genetic Improvement. Plants. 10.3390/plants13213073

47. Vincent, J., 1970. A Manual for the Practical Study of Root-Nodule Bacteria., IBP Handbo. ed. Blackwell Scientific Publications, Oxford, UK.

48. Wickham, H., 2016. ggplot2: Elegant Graphics for Data Analysis, Journal of the Royal Statistical Society Series A: Statistics in Society. Springer. 10.1111/j.1467-985x.2010.00676_9.x

49. Xin, P., Schier, J., Sefrenová, Y., Kulich, I., Dubrovsky, J.G., Vielle-Calzada, J.-P., Soukoup, A., 2022. The Arabidopsis TETRATRICOPEPTIDE-REPEAT THIOREDOXIN-LIKE (TTL) family members are involved in root system formation via their interaction with cytoskeleton and cell wall remodeling 946–965. 10.1111/tpj.15980

