## Supplementary figures and images for "Exploring the potential role of the *TETRATRICOPEPTIDE THIOREDOXIN-LIKE* gene family in nitrogen-fixing and water-restricted soybean plants"

### Figure S1_Sainz et al.

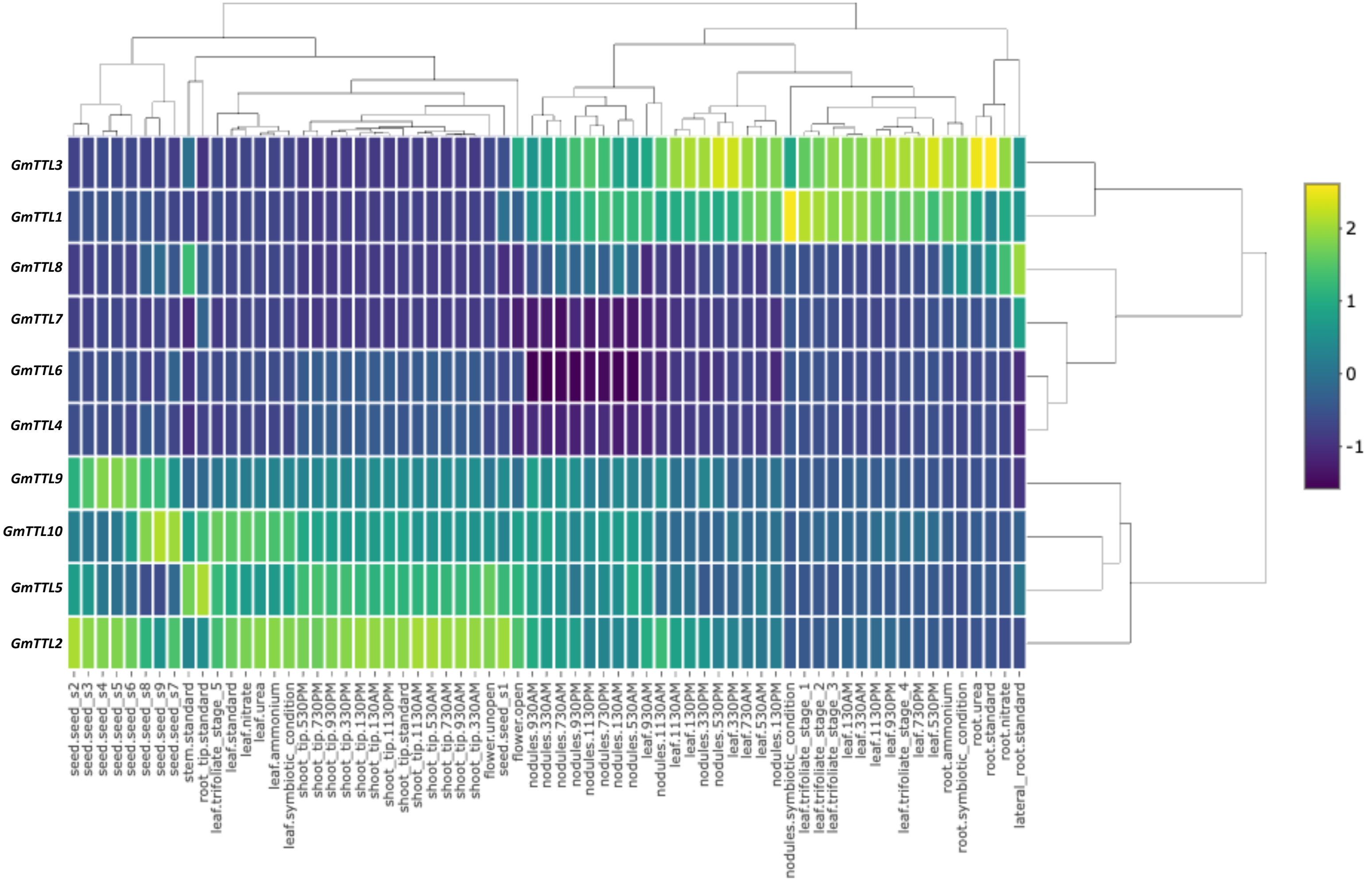
